# Conditional Diffusion with Locality-Aware Modal Alignment for Generating Diverse Protein Conformational Ensembles

**DOI:** 10.1101/2025.02.21.639488

**Authors:** Baoli Wang, Chenglin Wang, Jingyang Chen, Danlin Liu, Changzhi Sun, Jie Zhang, Kai Zhang, Honglin Li

## Abstract

Recent advances in AI have enabled the accurate prediction of a single stable protein structure solely based on its amino acid sequence. However, capturing the complete conformational landscape of a protein and its dynamic flexibility remains challenging. In this work, we developed Modal-aligned conditional Diffusion (Mac-Diff), a score based diffusion model for generating the conformational ensembles for unseen proteins. Central to Mac-Diff is an innovative attention module that enforces a delicate, locality-aware alignment between the conditional view (protein sequence) and the target view (residue pair geometry) to compute highly contextualized features for effective structural denoising. Furthermore, Mac-Diff leverages semantically rich sequence embedding from Protein Language Models like ESM-2 in enforcing the protein sequence condition that captures evolutionary, structural and functional information. This compensates for protein structural heterogeneity more effectively than embeddings from structure prediction models that are possibly biased to the dominant conformation. Mac-Diff showed promising results in generating realistic and diverse protein structures. It successfully recovered conformational distributions of fast folding proteins, captured multiple meta-stable conformations that were only observed in long MD simulation trajectories and efficiently predicted alternative conformations for allosteric proteins. We believe that Mac-Diff offers a useful tool to improve understanding of protein dynamics and structural variability, with broad implications for structural biology, drug discovery, and protein engineering.

## Introduction

Proteins are fundamental building blocks of life and play an integral role in cellular and biological processes. Many proteins possess inherent flexibility that enables them to function through the interconversion of different conformational states with varying energy levels. Such dynamic nature has profoundly determined protein functional repertoire in various contexts including ligand interactions, enzymatic reactions, and molecular evolutions. As a result, accurately generating the conformational ensembles for given protein sequences is vital for elucidating the mechanisms underlying their functionalities with wide applications in biological and medical sciences^1–4^.

In the past, experimental structure determination primarily focused on a single or at best a few discrete—static protein structures due to the significant cost^5^. In order to have a comprehensive delineation, molecular dynamics (MD) simulations are thus widely used to produce coherent trajectory of protein conformations^6^. Due to the high computational complexity and the requirement to simulate long timescales, MD simulations could be very time consuming and resource demanding^7–9^. The emergence of AlphaFold2 has dramatically advanced the state-of-the-art of protein structure prediction. By integrating structural and co-evolutionary information with Evoformer attention in an end-to-end learning architecture, AlphaFold2 allows faithfully predicting an individual (arguably, the most probable) protein structure^10^. However, as a deterministic sequence-to-structure mapping, it may not fully account for the protein conformational heterogeneity^11, 12^. A number of variants^13–19^ based on AlphaFold2 have emerged to produce multiple conformations for a protein by expanding the output of AlphaFold2. Examples include modifying multiple-sequence alignments (MSA) depth through subsampling^13, 14^, residue replacement^15^, MSA cluster level structure prediction^16^, using state specific structural templates^17^ and using multiple structural outputs as initialization for enhanced sampling^18^. Although with great potential, the generality of these techniques in predicting conformation heterogeneity or generating protein conformational distribution remain to be further explored^20, 21^.

In recent years, non-deterministic generative models drew considerable attention towards systematically generating the conformational ensembles of proteins. Early works focused on Generative Adversarial Networks or Variational Autoencoders (VAE)^22–24^. Then a significant amount of interest was drawn on diffusion models^25^ due to its promising results in generating novel and realistic samples from the distribution it is trained on. Examples include non-conditional diffusion models such as FoldingDiff^26^ and Str2Str^27^, and a number of conditional generative models using sequence representation from structure prediction models (as conditions) and various types of equivariant networks (for denoising), such as Eigenfold^28^, DiG^29^ and ConfDiff^30^, and those using advanced flow models and diffusion transformers such as AlphaFlow^31^ and IdpSAM^32^. Furthermore, diffusion models were also applied successfully in protein design tasks^33–36^.

Most existing conditional diffusion methods^27–30^ relied closely on structure prediction models such as AlphaFold2^10^, ESMFold^37^, or OmegaFold^38^ in terms of the protein geometric representation (e.g., residue frames^10^, *C_α_* atom coordinates^28^) and the denoising network architectures used in these methods (e.g., Invariant Point Attention^10, 39^). On the other hand, the initial sequence embedding was also obtained from these structure prediction models as the condition for generative models. For example, EigenFold and DiG used the residue embeddings from OmegaFold and AlphaFold2, respectively, and ConfDiff used the sequence embedding from ESMFold. Though encouraging results have been observed, exploration on whether sufficient structural heterogeneity could be extracted from sequence representations of structure prediction models remains to be continued. This is because most of these structure prediction models were originally designed to predict a single structure for a given sequence, and so the resultant representations might be possibly biased towards a dominant structure that structure-models tend to predict ^40^.

In this work, we present Modal-aligned conditional Diffusion (Mac-Diff), a score-based conditional diffusion model to generate realistic and diverse protein conformational ensembles. Mac-Diff performs iterative denoising on protein backbone geometries (target view) by continually receiving guidance from the protein sequence (conditional view). Here, instead of using structural prediction models like AlphaFold2, Mac-Diff adopts Protein Language Models (PLMs) like ESM-2^37^ to obtain the initial representation of protein sequences as conditions. ESM-2 was trained by unsupervised masked language modelling based on massive protein sequence datasets, allowing it to capture a wide spectrum of information ranging from evolutionary patterns, structural motifs, functional properties to broader biological knowledge at different scales. This semantically rich sequence representation enables Mac-diff to effectively compensate for the structural heterogeneity of proteins. Central to Mac-Diff is a new attention module to bridge the gap between the conditional modality (protein sequence) and the target modality (residue geometry) called Locality-Aware Modal Alignment attention (LAMA-attention). Compared with text-to-image tasks^41^ requiring only loose, unstructured alignment between text tokens and image pixels, LAMA- attention enforces a physically more delicate alignment between sequence and structure^1^, which restricts the attention field of each residue to only its local interacting environment. By this locality-aware alignment between the two views (or modalities), Mac-Diff was able to compute highly contextualized features in the target space to recover realistic and diverse protein structures. See Supplementary Note 1 for a comparison on the different levels of modal alignment for text-to-image generation and protein conformational ensemble generation, and discussions on the limitations of conventional cross-attention in the latter task.

Mac-Diff showed promising results in generating realistic conformational ensembles for given protein sequences. Empirically, we showed that Mac-Diff effectively recovered the conformational distribution of 12 fast folding proteins from the benchmark test set^27, 42^ it has never seen before, in terms of a number of important evaluation metrics such as the Jensen-Shannon divergence on *C_α_* -*C_α_* distance distribution, radius of gyration distribution, and *C_α_* -*C_α_* distance distribution on top-two TICA-components. Notably, the conformations generated by Mac-Diff exhibited greater diversity compared to competing methods, while maintaining a comparable level of accuracy in ensemble distributions. Furthermore, Mac-Diff showed encouraging performance on predicting alternative conformations for proteins that it has not seen before. For example, it recovered important conformational substates of BPTI that were observed in long MD simulations of 1 ms, and also predicted the closed state and the open state of Adenylate Kinase(AdK), an allosteric protein involved in energy metabolism. Finally, compared to traditional MD simulations, Mac-Diff achieved a sampling speedup of over 50 times. Overall, we believe that Mac-Diff has the potential to improve our understanding of protein folding dynamics and provide insights into the intricate relationship between protein sequence, structure, and function. The capability of Mac-Diff in predicting conformational heterogeneity will also be useful in applications of structure-based drug design and protein engineering.

## 1 Results

Fig. 1 illustrated the overall design of Mac-Diff. Fig. 1a is the backbone geometric representation used in Mac-Diff, including pairwise *C_β_* distance, dihedral angle, planar angle and a padding channel, which are invariant to 3D rotation and translation. Fig. 1b is the model overview. It is a score-based conditional generative model capable of recovering the conformational distribution of a protein by generating backbone geometric structures and converting them to atom-level coordinates through the Rosetta folding protocol. The forward diffusion process iteratively injects noise to geometric tensor, and the backward process achieves iterative denoising. The denoising network is a U-Net structure with five downsampling/upsampling stages. Each stage has a ResNet block to integrate time-step embedding and residue-pair representation, and a TransFormer block to update residue-pair representations through self-attention and LAMA-attention. Fig. 1c is the locality-aware modal alignment attention (LAMA- attention). It enforces a well-controlled spatial alignment between the sequence view and the structure view by forcing each residue to attend to only those neighboring residues with high contact probability, so as to update residue-pair representations with highly relevant and contextualized sequence features for denoising. More detailed descriptions can be found in the Methods section.

**Figure 1.**
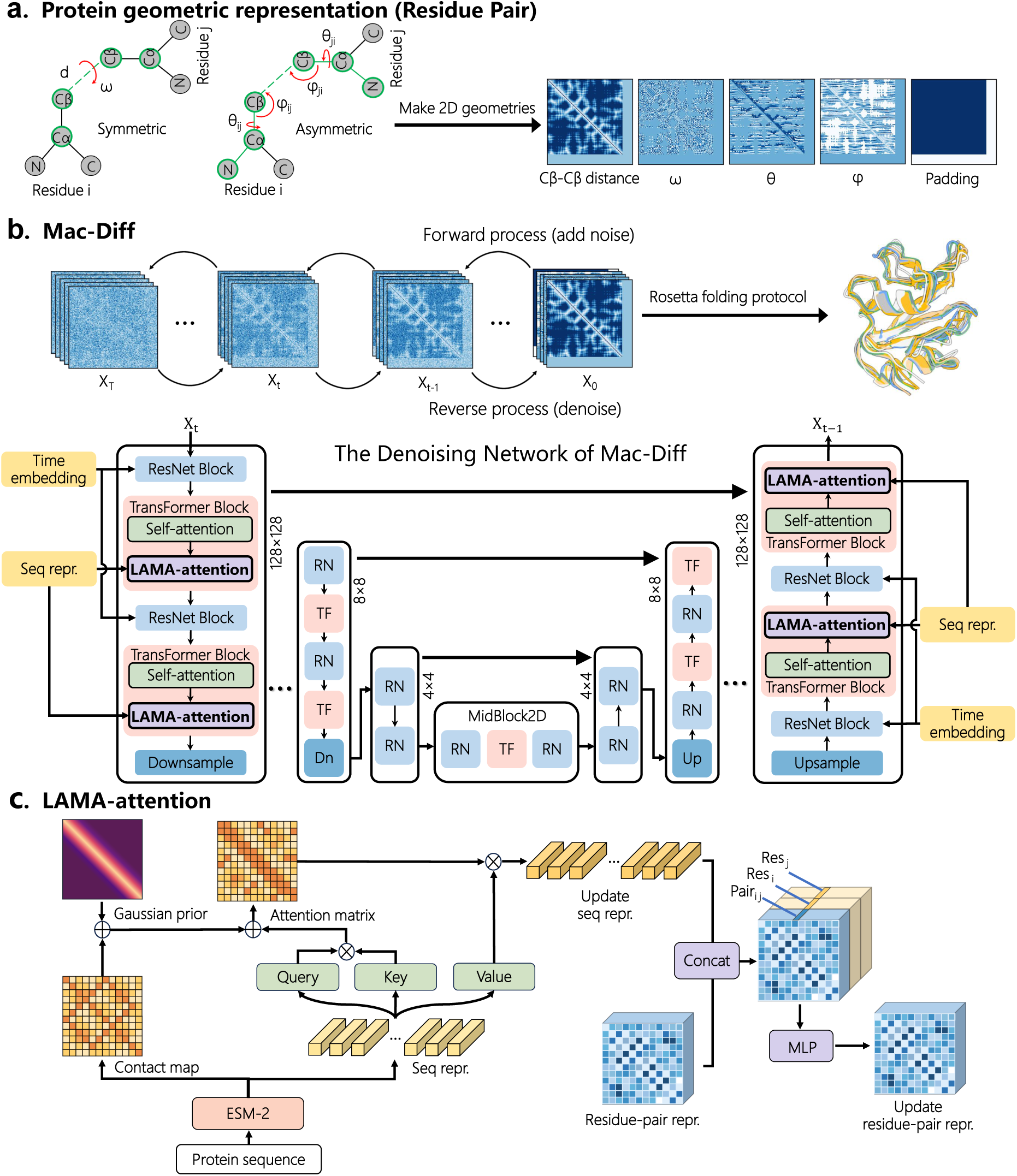
Overview of Mac-Diff architecture. **(a)** Protein backbone geometric representation with *L* residues as an *L×L×* 5 tensor with pairwise *C_β_* distance, dihedral angle *ω* along two *C_β_* atoms, dihedral angles *θ*, and bond angles *φ* (direction of *C_β_* atom of one residue in a reference frame centered on the other residue), and a padding channel indicating sequence length. **(b)** Mac-Diff workflow. The forward diffusion process iteratively injects noise to geometric tensor, and the backward process achieves iterative denoising.The denoising network is a U-Net structure with 5 downsampling/upsampling stages, each stage with ResNet block and TransFormer block (self-attention and LAMA-attention). **(c)** Locality-aware modal alignment attention (LAMA-attention), which allows each residue to attend to only those neighboring residues with high contact-probability, so as to update residue-pair representations with highly relevant and contextualized sequence features for denoising.

Fig. 2 illustrated the difference between Mac-Diff and stable diffusion - a most popular diffusion framework for text-to-image generation^41^, in their attention modules. In stable diffusion, the attention between pixels (query) and words (keys) was dense and global, indicating that the alignment between pixels and words were unstructured, purely data-driven, and were without prior algorithmic controls (Fig. 2a). In comparison, LAMA-attention allowed focusing only on the interacting neighbors (their amino acid types) of a residue when updating its representation (Fig. 2b). This effectively narrows the attention field from the whole sequence to only a small fraction of useful residues in the conditional view, marking a key distinction from the global cross-attention module in stable diffusion.

**Figure 2.**
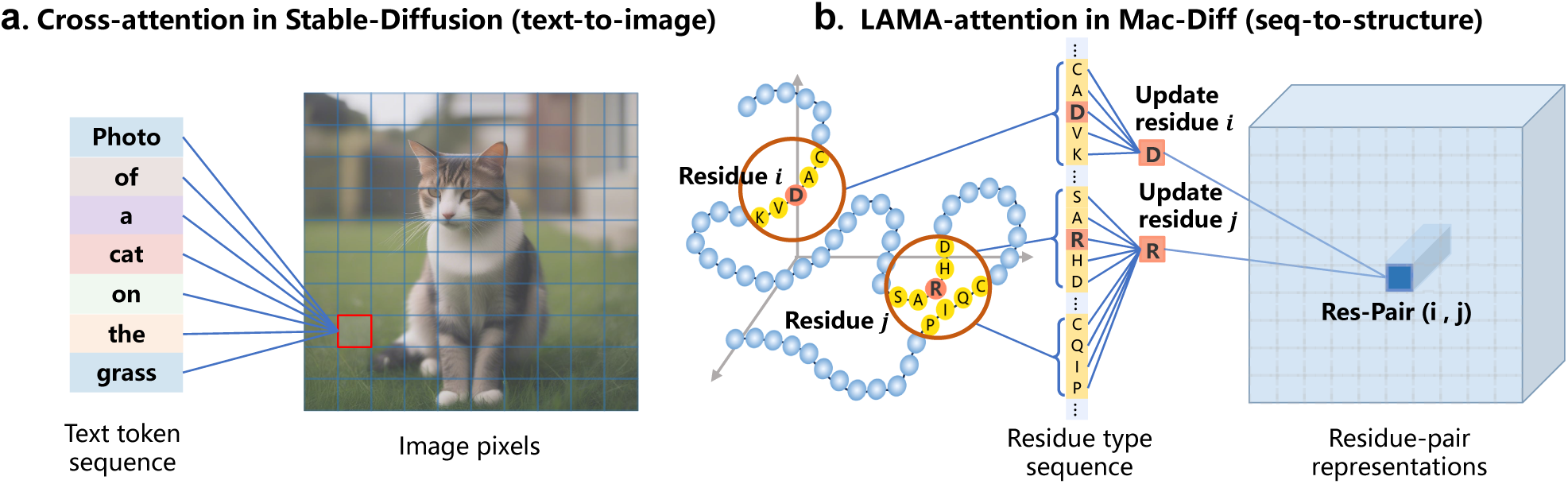
Schematic comparison of cross-attention (in text-to-image generation) and LAMA-attention (in protein conformational ensemble generation). **(a)** In traditional cross-attention, each pixel in the generated image is potentially related to all tokens in the input text without prior algorithmic control on the attention field. **(b)** In the locality-aware modal alignment attention (LAMA-attention), each residue-pair representation *i j* is related to only those residues that are likely to interact with residue *i* and *j* biologically - a stronger, locality-aware spatial alignment between sequence and structure.

We performed extensive evaluations of Mac-Diff against state-of-the-art methods for generating realistic and diverse protein conformational ensembles, using carefully curated training datasets and public benchmark testing datasets. Two types of tasks were performed: (1) Recovering the distribution of protein conformational ensembles and key conformational substates using target protein MD trajectory as reference (Sections 1.1 and 1.2), and (2) predicting alternative protein conformations using experimentally validated 3D protein structures as reference (Section 1.3). For task-I, we collected MD data from two public sources, GPCRmd and Atlas of proTein moLecular dynAmicS (ATLAS), leading to 1,674 protein trajectories as the training data and obtained the model “Mac-Diff-MD”, and tested its performance on the fast folding proteins and BPTI benchmarks^27, 42^; for task-II, we used the protein structures from the Protein Data Bank (PDB) as the training data and obtained model “Mac-Diff-PDB”, and tested its performance on a typical model protein AdK as well as 40 proteins from the Cfold dataset^43^.

### 1.1 Performance Evaluation on fast folding proteins benchmark

In this section, we evaluated Mac-Diff in recovering the conformational distribution of fast folding proteins. We trained our model using altogether 1,674 MD trajectories (371 from GPCRmd dataset^44^, and 1,303 non-membrane proteins from ATLAS dataset^45^), see detailed settings and statistics of the training data in the Training Data section. In the data preprocessing stage, we rigorously excluded the testing proteins from the training cohort, in order to strictly evaluate the generalization capacity of our model.

We used the MD training data for the fast folding protein and BPTI benchmark (task-I) because this task is primarily designed to assess the quality of the generated conformational distributions using MD simulations as ground truth. In comparison, PDB dataset has much more diverse conformations than short MD simulation could capture, and so it is more suited for task-II of predicting protein alternative conformations. In the literature, due to the scarcity of MD simulations, most generative methods used PDB data for training, and MD data were less used^27, 28, 30^. Our work thus represented a useful attempt to explore limited MD data in predicting protein conformational distributions.

We first evaluated Mac-Diff in generating conformational ensembles of fast folding proteins. Here, we chose 12 fast folding proteins including examples of all three major structural classes (*α*-helical, *β*-sheet, and mixed *α*/*β*) that were widely used as benchmark for evaluating the quality of equilibrium distributions generated by computational models^27, 30, 42^. The reference conformation distribution for each protein is represented by 1,000 conformations sampled with fixed stride from their MD simulation trajectories, ranging from 100 *µ*s to 1 ms to cover multiple folding or unfolding events^46^. See detailed protein trajectory information in Section 3.1.2 and Supplementary Table 6.

We then generated 1,000 conformations using Mac-Diff and each of the competing methods to recover the protein conformational distribution, and examined the quality by assessing their deviation from the reference MD distribution (as ground truth). To provide a quantitative analysis, we computed the Jensen-Shannon divergence on the following distributions, including: (1) pairwise *C_α_* atom distance distribution (JS-PwD), which reflects the global spatial structure of the protein; (2) radius of gyration distribution (JS-Rg), which is based on the distance between each *C_α_* atoms to the center of mass of the protein, reflecting the compactness of the protein, and (3) pairwise *C_α_* atoms distance distribution computed in the top-2 time-lagged independent components (JS-TIC), which is commonly employed to analyze slow dynamics of proteins MD trajectories^47–49^. The metrics employed to quantify the generated conformational distributions are evaluated independently for each protein and subsequently averaged across the 12 fast folding proteins. Detailed definitions of the three metrics can be found in Section 3.1.3.

As shown in Fig. 3a, Mac-Diff showed competitive results when compared with existing diffusion-based and flow-based models across all equilibrium metrics on fast folding proteins. For the three error metrics averaged over the 12 fast folding proteins, i.e., JS-PwD, JS-Rg, and JS-TIC, Mac-Diff was about relatively 13%, 4%, 10% lower than the best competing method, respectively. In Supplementary Tables 1-3, we provided more detailed comparative results on each of the 12 fast folding proteins under these three metrics. These results showed the potential of Mac-Diff in recovering protein conformational ensembles.

**Figure 3.**
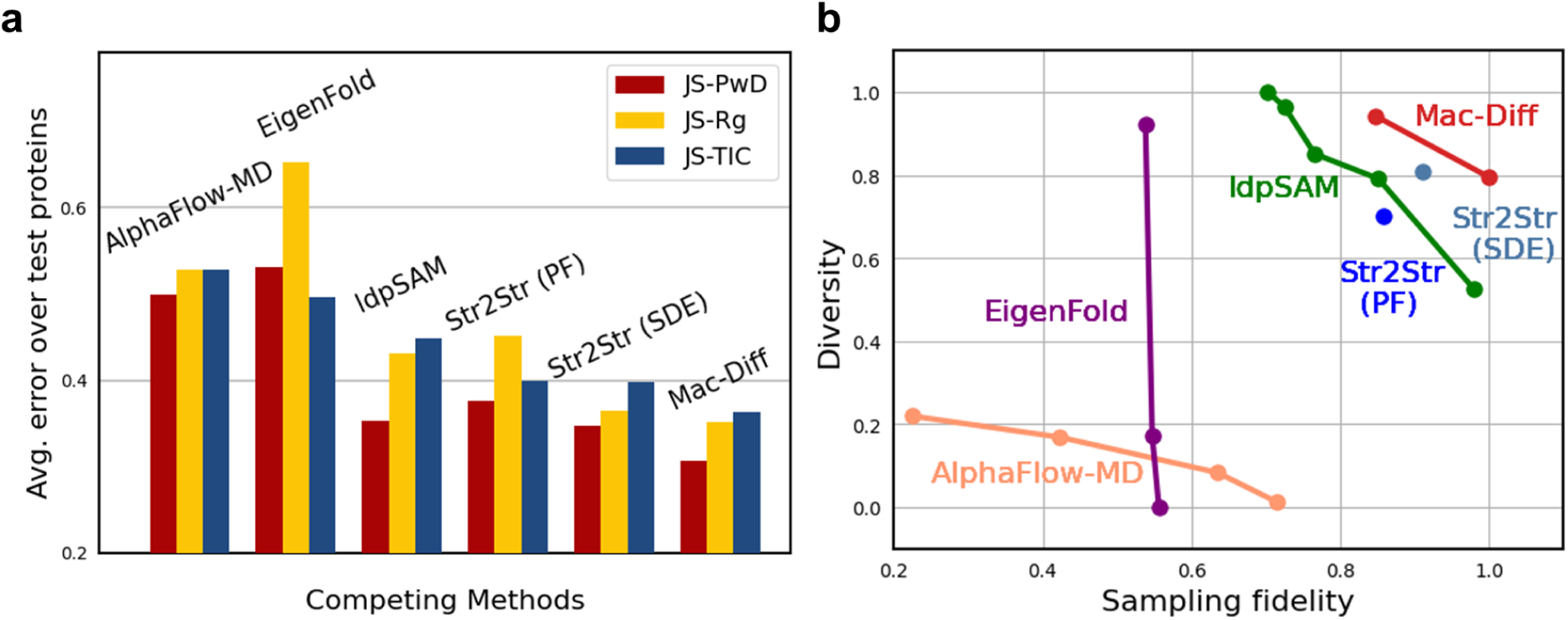
Performance of all competing methods in recovering the conformational distributions of the fast-folding proteins. **(a)** The Jason-Shannon divergence between the generated conformational distribution and the reference MD distribution for 6 competing methods averaged over 12 fast folding proteins^46^, when considering the pairwise *C_α_* atom distances (JS-PwD), the radius of gyration (JS-Rg), and the pairwise *C_α_* atom distances along the top-2 time-lagged independent components (JS-TIC). **(b)** The Pareto frontiers for each of the six competing methods, evaluated on the 12 fast-folding proteins in the test set, plotted in the *diversity-versus-fidelity* space. Higher values indicate better performance for both metrics. Each point represents the diversity-fidelity result for an individual protein, with only points on the Pareto frontier for each method shown. Points closer to the upper-right corner reflect better performance (trade-off between diversity and fidelity). Diversity values were normalized to [0, 1].

Next, we examined the diversity of conformations generated by different methods. Following the literature^30^, conformational diversity was quantified by the average pairwise distance among the generated conformations. However, evaluating diversity alone is insufficient. So we jointly assessed both the diversity of the conformation set and the fidelity of the conformational distribution measured by 1 *−JS_PwD_*, with *JS_PwD_* the Jensen-Shannon divergence between the recovered distribution and the reference MD distribution, as defined in equation (1). For both the diversity and fidelity metrics, higher values generally indicate better performance. In Fig. 3b, we presented the Pareto frontier for all competing methods evaluated on the 12 fast folding proteins, plotted in the diversity-versus-fidelity space. Points closer to the upper-right corner of this plane indicate better performance, as they reflect an optimal balance between the two metrics considered. As shown, Mac-Diff consistently achieved superior performance compared to all other methods, demonstrating its ability to generate protein conformations that are both highly diverse and faithful to the reference distribution.

We also used the time-lagged independent components analysis (TICA) to better visualize the conformational distributions generated from different models in a low-dimensional subspace. TICA is a dimension reduction technique for complex dynamic systems commonly applied in the study of protein folding and unfolding processes^47^. Following the practice of Lu et al.^27^, we have computed the top-2 TICA components from the pairwise *C_α_* atom distance matrix that reflects the slow and dominant transitions among different protein conformational ensembles. Fig. 4a visualized the conformations produced by MD simulation and six generative models for *α*3D, Protein B and Homeodomain, in which each conformation generated was projected (as a dot) to the top-2 TICA-components^47, 48^. The JS-TIC error metric for each generative model was also shown in the subfigures. We observed that the conformations generated by Mac-Diff could better approach the reference MD data, both visually and in terms of the JS-TIC metric. The JS-TIC by Mac-Diff was relatively 38%, 24%, 64% lower than the best competing method for the protein *α*3D, Protein B and Homeodomain, respectively. The JS-TIC error metrics for the remaining nine fast folding proteins, including Villin, BBA, Protein G, NTL9, WW domain, Lambda, Trp-cage, BBL and Chignolin were shown in Supplementary Table 3.

**Figure 4.**
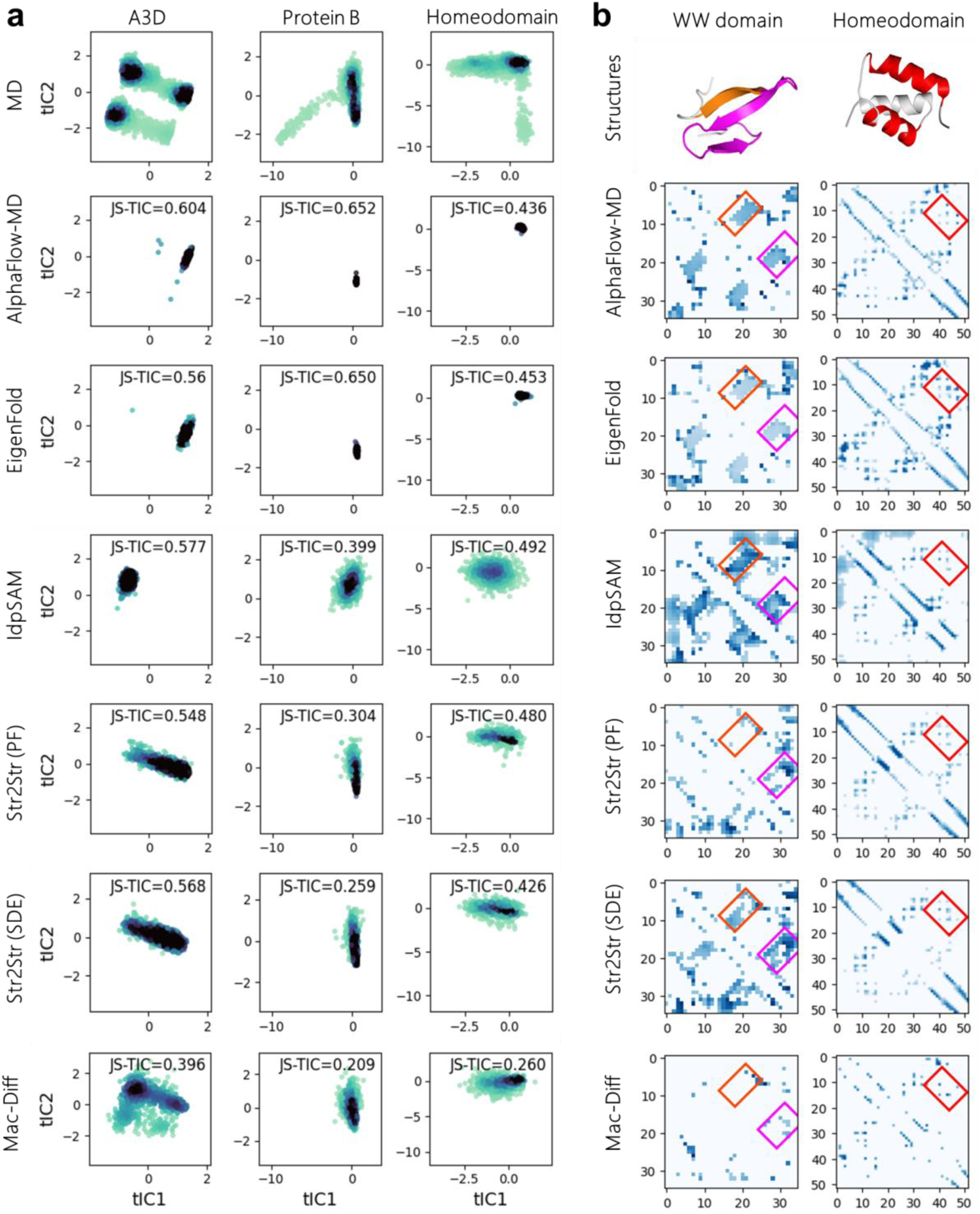
Performance of all competing methods in recovering protein conformational distributions and residue contact probabilities. **(a)** Conformations produced from MD-simulation and 6 generative models when projected onto the top-2 TICA-components for *α*3D, Protein B and Homeodomain. Each dot represents a conformation and is color coded by its density (the darker the point, the higher the density). **(b)** Approximation errors of the residue contact probability map (difference with the ground truth probability map) from 6 competing methods on WW domain and Homeodomain. The sparser/lighter the pixels, the lower the error. Contact patterns of the *β*-sheets in WW domain (boxed with orange and magenta) and the *α*-helix-*α*-helix interaction in Homeodomain (boxed with red) were shown in the upper triangles. The folding structures in the top were marked correspondingly by the same colors.

Fig. 4b presented the probability-map for pairwise residue contact relationships in two proteins, WW domain (with a three-stranded antiparallel *β*-sheet in folded structure), and Homeodomain (with three *α*-helices in folded structure), as recovered by all competing models. Here, the *i j*th entry in the probability map represents the probability of contact between the *i*-th and *j*-th residues across all generated conformations, using a threshold of 10 Å^42^. To better visualize the quality of the probability map recovered by each method, we plotted the difference between true/recovered probability map. We observed that Mac-Diff learned both local contact patterns and global structural patterns along the sequence^50, 51^. AlphaFlow-MD and EigenFold tended to predict stable native structures as opposed to those contact patterns arising from unfolded, but physically feasible structures. IdpSAM produced contact patterns that are more like in disordered proteins (e.g., *β*-sheet related contact patterns for protein WW domain marked in magenta box in Fig. 4b), which might be due to the fact that it was trained primarily on intrinsically disordered proteins. Str2Str tended to emphasize certain weak contact patterns between the second *β*-sheet and the C-terminus residues (boxed in magenta in Fig. 4b). See more visualization results in Supplementary Fig. 2. In Supplementary Table 4, we reported the root mean square error (RMSE) of the residue contact probability map for all the competing methods, in which Mac-Diff had the lowest average approximation error across the 12 fast folding proteins. This illustrated the effectiveness of Mac-Diff in faithfully capturing key contact patterns in protein folding processes.

Homeodomain is a class of evolutionarily conserved protein with three *α*-helices. Commonly observed 3D structures are compact with a low JS-Rg value (average distance between *C_α_* atoms and protein centroid which can be deemed as the radius of the protein), in which the first and the last helices form intra-chain contacts due to hydrogen bonding of side chain atoms. We carefully examined the contact-map recovered by Mac-Diff, in particular its sub-block corresponding to the residues from the 1st and the last helices. As can be seen in Fig. 4b and Supplementary Fig. 2, Mac-Diff can accurately capture contact probabilities, leading to a spatially compact 3D structure that is well consistent with experimental observations.

### 1.2 Conformational substates prediction for the BPTI

To further examine the capacity of Mac-Diff in capturing protein conformational changes, we used BPTI, a globular protein of 58 residues, for a case study. In the literature^52^, Shaw et al. found that BPTI had five structurally distinct representation clusters in a long MD simulation of 1 ms. Here, in Fig. 5, we visualized the representative conformation for each of these five clusters, as well as the conformations generated from all the competing methods, by embedding them in the top-2 time-lagged independent components (TICA-components) of the reference MD trajectories^27^, corresponding to slowly varying molecular dynamics that are of the most interest^47–49^. We also plotted the contours of the 2D-density of MD simulated conformations by a kernel density estimator with bandwidth determined by the default Scott’s Rule^53^.

**Figure 5.**
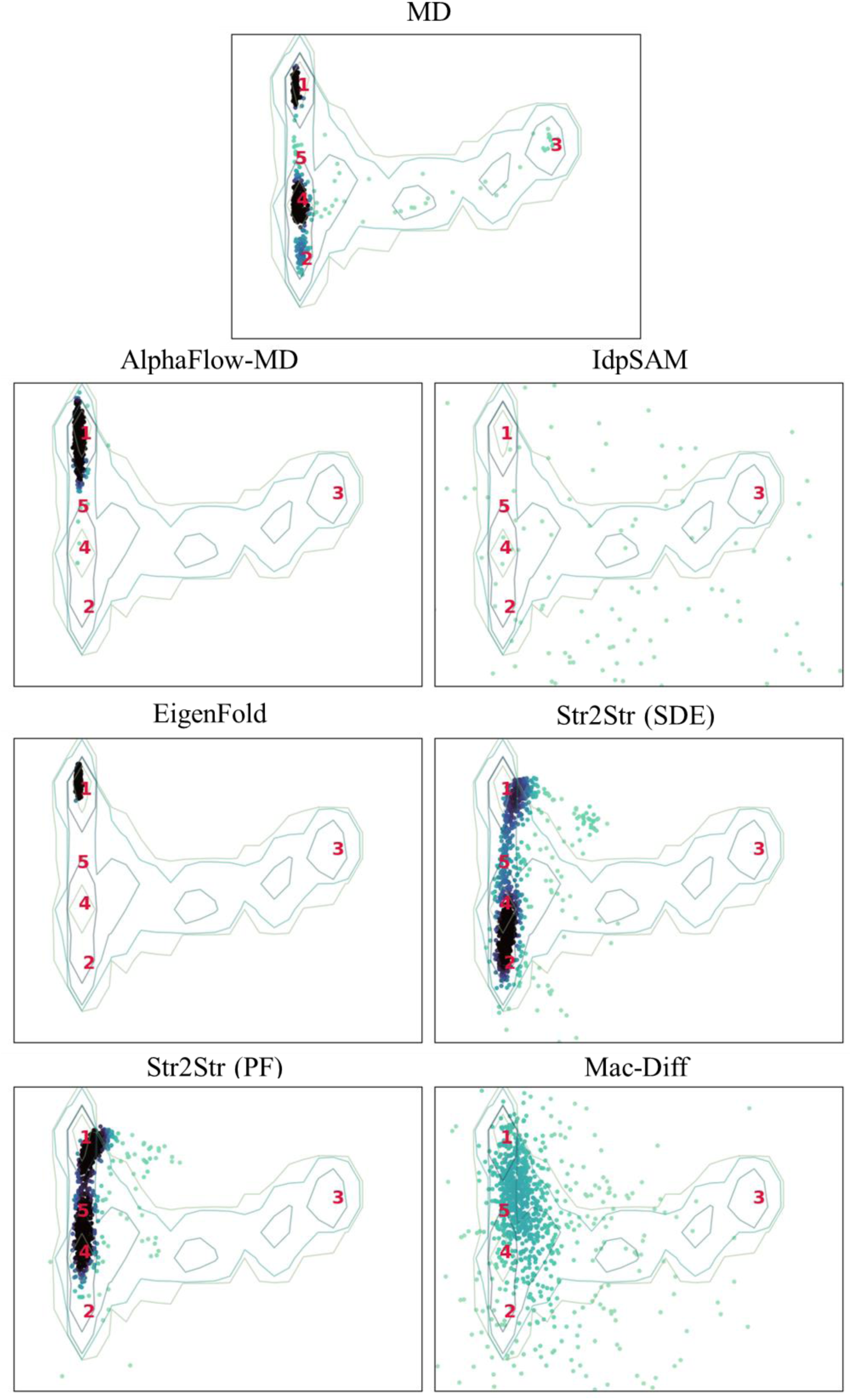
Generated conformations (dots) from six competing methods, along with the reference MD distribution for BPTI, visualized in the top two TICA-components most relevant to protein folding dynamics. The density contours of the MD reference distribution were plotted in each result. Five key conformation clusters were marked by numbers 1-5 in red color.

In Fig. 5, we observed that for EigenFold and AlphaFlow-MD, most of the generated conformations were centered around cluster 1. In fact, capturing protein conformations across long time scales poses a significant challenge. The conformations generated by IdpSAM were relatively scattered, indicating that it might be more suited for intrinsically disordered protein’s (based on which IdpSAM was trained^32^). For Str2Str (SDE) and Str2Str (PF), the generated conformations well covered four conformational substates but substate 3, which is separated from the other states by relatively steep energy barriers on the energy landscape. In comparison, Mac-Diff generated samples that were closer to substate 3, in addition to samples around the other four clusters. Quantitatively, in terms of the RMSD between the generated conformation and the representative conformation of each cluster (subject to a translation-and-rotation based calibration between them) as used in the literature^27^, Mac-Diff had the lowest RMSD for cluster-3, and ranked either the 2nd or the 3rd best among all the six competing methods for clusters 1, 2, 4 and 5. See Supplementary Table 5 for the RMSD errors of the six competing methods for all the five distinct conformation clusters.

In our experiments, we rigorously excluded all BPTI related MD trajectory data in training the Mac-Diff model, thus guaranteeing the validity of the evaluation metrics reported. For the other 5 competing methods, however, since we directly downloaded their models in our evaluations, we did not have any control of their training data, but simply reported their results for a complete comparison.

In order to verify the validity of the generated conformational ensembles of the BPTI by Mac-Diff, we further plotted the bond length distribution and backbone dihedral angle distribution in Supplementary Fig. 6a and b. We observed that Mac-Diff achieved a good agreement with the MD reference data, in that the mean bond length of the 1,000 conformational structures, 3.81 Å, is very close to the mean bond length of reference MD (3.85 Å) and the ideal bond length (Previous study on protein structure quality assessment in PDB demonstrate that the distance between consecutive *C_α_* atoms are distributed normally with a mean of 3.8 Å^54^). Moreover, the distribution of the distances between non-bonded *C_α_* atoms that are at least four residues apart in the protein sequence were also shown in Supplementary Fig. 6a (right). The lower bound of the non-bonded atom distances by Mac-Diff is 3.49 Å, which is very close to that derived from the reference MD (3.48 Å). This indicated that the backbone self-crossings in the generated conformational ensembles by Mac-Diff were successfully avoided.

In Supplementary Fig. 6b, we evaluated the torsion angles *φ* and *ψ* generated by Mac-Diff in terms of the Ramachandran map^55^, which describes the key rotational freedom around backbone peptide bonds. Results showed that most of the generated angle pairs fell into the region of right-handed *α*-helix conformations, *β*-sheet conformations and left-handed *α*-helix conformations, closely matching the angles distribution of the reference MD trajectories. Taken together, these results showed that Mac-Diff was capable of generating valid conformations and capturing important substates with moderate diversity and quality when compared to a series of diffusion-based and flow-based generative models.

### 1.3 Capturing alternative conformations of allosteric proteins

In this section, we investigated the capacity of Mac-Diff in capturing multiple important conformational states of a model protein AdK in Escherichia coli, and proteins in the Cfold40 test set. The Cfold40 contains 40 proteins selected from Cfold^43^ dataset such that (1) each protein has less than 256 residues (out of computational cost considerations), and (2) each protein has two alternative conformations with the TM-score (a metric for assessing the topological similarity of two structures)^56^ between the two conformations below 0.8. See more details in Section 3.1.2. Note that capturing multiple conformational states of a protein is crucial to unrevealing its functional aspects that a single structure may fail to reveal.

In this task, we trained Mac-Diff using PDB structure data (Mac-Diff-PDB) as many other generative models in the literature^28, 29^, because PDB dataset has much greater sequence diversity and conformational variations than our MD data used in the previous tasks (short simulations up to 500 ns for 1,674 proteins). To insure a legitimate evaluation, we removed those proteins from the training data whose sequences had a significant similarity with those of the testing proteins (sequence alignment scores larger than 90% by Ch-HIT^57^). See detailed Mac-Diff-PDB model training protocols in Section 3.3. In Supplementary Table 7, we provided a detailed summary of Mac-Diff and other competing methods in terms of how they chose the training data from PDB, and how they obtained reside embedding vectors.

We have included 4 state-of-the-art generative models in our comparisons, including (1) DiG^29^, a diffusion model trained on both PDB and more than one thousand MD trajectories with initial residue embeddings from AlphaFold2’s Evoformer; (2) MSA subsampling^13^, an AlphaFold2-based method generating protein alternative conformations by performing MSA subsampling; (3) AF-Cluster^16^, an AlphaFold2-based method generating multiple conformations by MSA clustering; (4) AlphaFlow-MD^31^, which fine-tuned the weights of AlphaFold2 on OpenProteinSet and MD datasets under flow matching framework. Note that the DiG model needed to use Evoformer’s embedding from AlphaFold2 as part of its input, but only released the embedding files of a few test proteins. In our evaluations, we thus picked one protein, AdK from their release, based on which we made comparisons with DiG; For the other 3 competing methods (MSA subsampling, AF-cluster and AlphaFlow), since they did not request external feature files, we were able to make a full comparison with them on all the 40 proteins in Cfold40 test set. For these three competing methods, we did not report their results on AdK because the two conformational states to be predicted appeared to be included in their training data.

We first compared the sampling capability of Mac-Diff-PDB and DiG on the protein AdK. Fig. 6a and b showed their sampling results. The native structure of AdK contains three key domains: a relatively rigid CORE domain, and two highly flexible domains, the NMP and LID domains, which are known to undergo conformational transitions between open and closed states during the adenosine triphosphate (ATP) to adenosine diphosphate (ADP) catalysis^58–60^. We observed that Mac-Diff-PDB exhibited a high conformational diversity covering both of the open state and the closed state: the highest TM-score between the generated structures to the two representative states was 0.85 and 0.90, respectively. In comparison, the sampling results of DiG were more centered around the closed conformation state corresponding to a low-energy state. It’s worthwhile to note that when allowed to perform a larger number of sampling (about tens of thousands)^29^, DiG could recover both conformational states of AdK.

**Figure 6.**
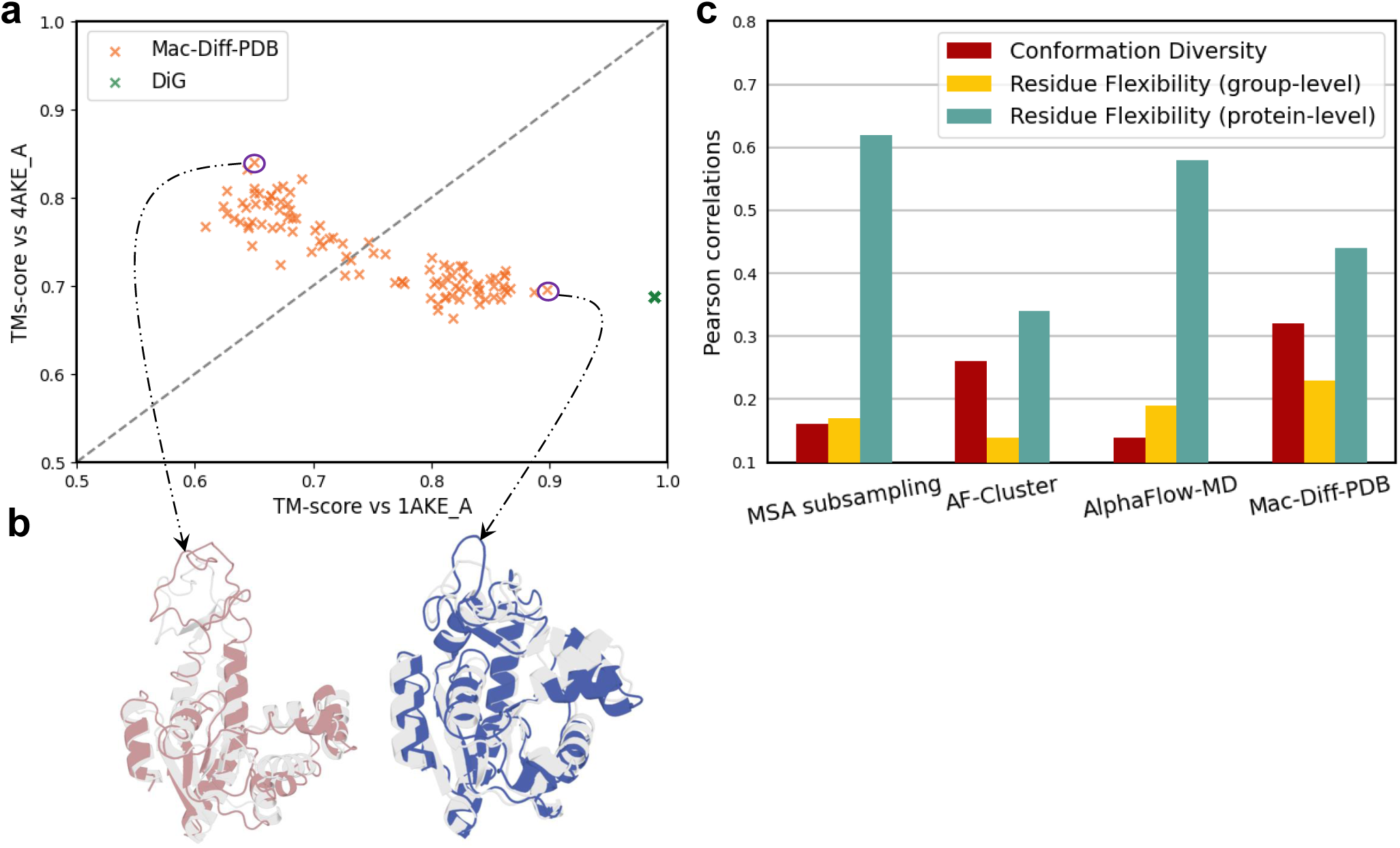
Sampling performance of Mac-Diff-PDB and competing methods on the allosteric protein AdK and the Cfold40 test set. **(a)** Scatter plot of TM-scores measure the structural similarity between generated 100 conformations and two experimentally determined states: closed (PDB ID: 1AKE_A) and open (PDB ID: 4AKE_A). **(b)** Overlay of Mac-Diff-PDB generated structure (blue/claret) having the best TM-score with the ground-truth conformations 1AKE_A/4AKE_A (gray). **(c)** Performance of Mac-Diff-PDB and three other generative methods based on the Pearson correlations between the generated conformations of a protein and its two ground-truth structures in terms of conformation diversity, residue flexibility at group-level and protein-level, respectively. The correlations are computed based on all the proteins in Cfold40 test set, see Section 3.1.3 for detailed definitions.

Next, we reported the comparisons between Mac-Diff-PDB and other 3 competing methods i.e., MSA subsampling^13^, AF-cluster^16^, and AlphaFlow^31^ on the 40 proteins in the test benchmark Cfold40. All methods were used to generate 100 conformations for every protein, and all MSAs used in this evaluation were fetched from the ColabFold server^61^. The detailed settings for these methods were as follows. (1) For AF-Cluster, the first 100 MSA clusters generated were selected as input, and then a local AlphaFold2 (version2.3.1) was executed for each MSA cluster to produce the final protein structure conformations with default parameters. (2) For MSA subsampling, we used 5 MSAs with depths 32, 64, 128, 256 and 1024; ‘max_msa_clusters’ was set to half the MSA depths^13^, except MSA-depth 1024, for which ‘max_msa_clusters’ was set to 128 following the setting of AlphaFold2. Following MSA-subsampling^13^, we used all the five weight files released by AlphaFold2, 4 random seeds, and a recycling number of 1 to generate altogether 100 conformations for each protein (4 seeds by 5 MSAs by 5 weight files). (3) For AlphaFlow, we used AlphaFlow-MD model ^31^ to generate 100 conformations for each protein with the default parameters. (4) For Mac-Diff-PDB, to reduce the inference cost, we trained two versions in charge of sequences with length range [1,128] and [128, 256], respectively. The ESM2-650M and ESM2-3B were used for Mac-Diff-PDB in computing the residue embedding.

Fig. 6c showed the performance of different methods in terms of how well they approached the ground-truth in terms of protein structural diversity. Here we have used the Pearson correlation (between estimate / ground truth) for two metrics: (1) Conformation diversity (TM*_ens_*), which describes (pairwise) diversity among different conformations of a protein; (2) Residue flexibility, which describes the spatial flexibility of a residue across different, but globally aligned conformations of a protein. In particular, following Jing et al.^28^, we computed the correlations of residue-flexibility at both group-level (all residues across all proteins in Cfold40) and protein level (residues for each protein in Cfold40, with mean and median correlation reported). See detailed definitions of these metrics in Section 3.1.3.

As can be seen in Fig. 6c, in terms of the conformation diversity, Mac-Diff-PDB had the highest correlation with the ground-truth, which is 0.32. Supplementary Fig. 7 (lower) further presented the scatter plot of the estimated conformation diversity by Mac-Diff-PDB (TM*_var_*) versus the true diversity (TM*_con f_* _1_*_/con f_* _2_) across all the 40 proteins in the Cfold40 test set. This showed that Mac-Diff-PDB could better preserve the pairwise diversity of protein conformational ensembles relative to the true structural ensemble. In terms of the group-level residue flexibility, Mac-Diff-PDB also showed the highest correlation, which is 0.23, meaning that the residue-level variations within the predicted samples were more predictive of true residue flexibility under the conformational change compared with other models. In terms of the protein-level residue flexibility, Mac-Diff-PDB was relatively lower than MSA-subsampling and AlphaFlow-MD. The discrepancy between group-level and protein-level flexibility estimation performance suggests that our method may be more susceptible to aggregation bias, particularly for proteins with markedly different flexibility profiles. In future studies, we plan to explore strategies to enhance the adaptability of Mac-Diff in accommodating proteins with diverse structural variability.

Supplementary Fig. 7 (upper) presented the ensemble TM-score (TM*_ens_*) versus the ground-truth conformation diversity (TM*_con f_* _1_*_/con f_* _2_) for all the proteins in the Cfold40 test set. The TM-ensemble score measures how well the generated structures for a protein cover its ground-truth structures (two in our studies). Here, we found that MSA subsampling, AF-cluster, AlphaFlow-MD demonstrated comparable results, surpassing the performance of the baseline (a hypothetical model which perfectly predicts one of the two conformations^28^, marked by a line with a slope of 0.5 and an intercept of 0.5) for 21, 18, and 16 cases out of the 40 proteins in Cfold40 test set. In comparison, Mac-Diff-PDB exhibited lower performance. Note that Mac-Diff-PDB used the protein language model ESM-2 for generating sequence representations. In comparison, the competing methods leveraged the Evoformer module of AlphaFold2 in constructing initial sequence representations, which we speculate can be more advantageous in producing high-quality protein folding structures. In the future, we will continue to improve the structural prediction capacity of Mac-Diff with advanced diffusion architecture^62, 63^ and incorporate the Rosetta folding protocols into a completely differentiable framework^64^.

In Supplementary Note 6 and Supplementary Fig. 8, we further analyzed and visualized five cases from the Cfold40 test proteins, where the TM-scores between the best conformations generated by Mac-Diff-PDB and the two ground truth structures both exceeded 0.8.

### 1.4 Mac-Diff sampling speed

In this section, we compared the sampling speed of Mac-Diff with MD simulations on a single NVIDIA A100 GPU. As shown in Supplementary Table 8, when compared to traditional MD simulations executed on OpenMM^65^ (see detailed settings in Section 3.4). Mac-Diff offers a more efficient alternative with a speed about 50 times faster. The performance is superior to 100 *µ*s MD simulation in terms of the JS-PwD and JS-Rg metric, although it is slightly worse than 100 *µ*s MD simulation in terms of the JS-TIC metric. It is worth noting that the inference speed of Mac-Diff is still slower than other diffusion based model (e.g., Str2Str needed about 10 minutes), because we adopted a naive implementation of diffusion process with about 2,000 denoising steps. Importantly, this process can be drastically accelerated by exploiting fast sampling methods or SDE solvers for diffusion models in the literature^66–68^, which can produce high-quality samples with up to 5-10 times or more reduction in the sampling time steps requested. We will continue to work on more efficient inference procedures for Mac-Diff in our future studies.

## 2 Discussion

We presented Mac-Diff, a conditional diffusion model to recover protein conformational ensembles. Mac-Diff is characterized by an innovative, locality-aware attention module that aligned the residue neighborhood across the conditional and the target views spatially in a more stringent manner. In particular, the interacting neighbors of each residue, which is key to acquiring structural representations for effective structure denoising, were determined carefully via the combination of three sources, namely, an isotropic Gaussian kernel emphasizing local residues along the 1-D chain structure, the contact matrix in the ESM-2 protein language model that was acquired (almost) in an unsupervised manner through pre-training with tens of millions of protein sequences, and residue interactions based on their embedding features. This comprehensive information allowed the model to control the alignment between the conditional and the target views precisely and in a locality-aware manner. Empirically, the LAMA-attention based diffusion showed promising results in recovering protein conformational distribution and identifying key conformational substates on fast folding proteins and BPTI benchmark. Furthermore, Mac-Diff successfully predicted alternative structures for allosteric proteins AdK even without relying on MSA.

There are a number of directions that we plan to study in the future. First, Mac-Diff was trained on only two publicly available MD databases with scanty time scales (from 100 *ns* to 500 *ns*) and limited protein diversity (1,674 proteins). The short MD simulations on these proteins may capture only small structural fluctuations, making it challenging to make predictions on larger proteins with higher degrees of structural freedom. Therefore, we plan to collect more MD simulation data with larger timescale (*µ*s or ms) scattered in the scientific literature, and curate them into large MD trajectories databases with unified and standardized simulation protocols. Second, although Mac-Diff was able to model protein conformational diversity, the flexibility at the residue level across proteins with different flexibility profiles remained to be improved. Combining generative models with MSA information, such as the sequence representation from the MSA Transformer^69^, was a direction worth studying in the future. Third, following the success of trRosetta and ProteinSGM in protein structure prediction and protein design, we will investigate the combination of the diffusion model and the energy minimization protocols in a complete end-to-end optimization framework^35, 64^ to improve the fidelity of the conformations generated. Furthermore, to improve the quality and speed of Mac-Diff generated conformations, we shall be considering more compact protein structure representations such as SE(3) backbone^34, 39^, and will employ advanced diffusion samplers such as EDM^70^. Finally, we plan to study generative model that can efficiently generate samples that conform to the Boltzmann distribution like the Boltzmann generator^71^, by utilizing a normalizing flow to model and generate equilibrium distributions for a specific molecular system.

## Acknowledgements

We thank Prof. Changlin Tian for the data collection and helpful discussions. This work was supported in part by the National Key Research and Development Program of China (2022YFC3400501); the National Natural Science Foundation of China (82425104, 62276099, 82173690).

## Author Contributions Statement

K.Z. and H.L. conceived the concept and designed the workflow for this study. K.Z. and J.Z. designed the locality-aware modal alignment attention module. B.W., C.S., C.W. and J.C. designed the diffusion model. B.W., C.W. and J.C. implemented the Mac-Diff network architecture. B.W., C.W., J.C. and D.L. performed computational experiments. B.W. and D.L. prepared all data and contributed to the analysis and interpretation of results. B.W., K.Z., D.L., C.S. and J.Z. assisted with designing the experiments. B.W., K.Z., C.W., D.L., J.Z. and H.L. wrote the manuscript text and prepared figures and tables. All authors provided critical feedback and helped shape the research, analysis, and revised the manuscript.

## Additional Information

**Supplementary Information** The supplementary material can be found at supplementary.pdf.

## Footnotes

1 Each residue in the sequence relates exactly to 3D coordinates of this residue in 3D space, but this “alignment” is trivial; it is the alignment between a residue in the 3D-space and its spatially interacting neighbors traced back to positions of the input sequence that are the key to injecting useful sequence information into the target space to generate denoised structures.

